# The ribbon architecture of the Golgi apparatus is not restricted to vertebrates

**DOI:** 10.1101/2021.09.10.459720

**Authors:** Giovanna Benvenuto, Maria Ina Arnone, Francesco Ferraro

## Abstract

The Golgi apparatus plays a central role as a processing and sorting station along the secretory pathway. In multicellular organisms, this organelle displays two structural organizations, whereby its functional subunits, the mini-stacks, are either dispersed throughout the cell or linked into a centralized structure, called Golgi “ribbon”. The Golgi ribbon is considered to be a feature typical of vertebrate cells. Here we report that this is not the case. We show that sea urchin embryonic cells assemble Golgi ribbons during early development. Sea urchins are deuterostomes, the bilaterian animal clade to which chordates, and thus vertebrates, also belong.

Far from being a structural innovation of vertebrates, the Golgi ribbon therefore appears to be an ancient cellular feature evolved before the split between echinoderms and chordates. Evolutionary conservation of the ribbon architecture surmises that it must play fundamental roles in the biology of deuterostomes.

## Introduction

In April 1898, Camillo Golgi reported to the Medical–Surgical Society of Pavia, his discovery of an intracellular structure in the neurons of the barn owl *Tytus alba*. In Golgi’s words, this structure appeared as “a fine and elegant network within the cell body … completely internal in the nerve cells … The distinctive appearance of this *internal reticular apparatus* is attributable to the prevalence of *ribbon-like threads*, their manner of dividing, their anastomoses, and the pathways formed by them…” (italics are ours)^1^. Golgi’s “internal reticular apparatus” later became known as Golgi apparatus or complex. With the advent of molecular tools and the transformation of classical cytology into experimental cell biology, the Golgi complex was shown to be central in the processing and sorting of secretory cargos^2,3^. The ribbon organization of the Golgi apparatus is believed to be typical of vertebrates, since in other multicellular organisms, such as flies, worms and plants, Golgi mini-stacks are dispersed throughout the cell cytoplasm^4-7^. The functional importance of the Golgi ribbon remains elusive. Here, we report that the Golgi ribbon is present in sea urchins. This observation indicates that whatever functions the Golgi ribbon might play they are not specific to vertebrate biology.

## Results

Published evidence is suggestive that in sea urchin embryos the Golgi complex can acquire a centralized morphology reminiscent of the ribbon structure observed in vertebrate cells^8,9^. These studies prompted us to analyse Golgi dynamics in the developing embryos of the Mediterranean sea urchin, *Paracentrotus lividus*.

Immediately after fertilization, *P. lividus* zygotes were microinjected with *in vitro* transcribed mRNAs encoding fluorescent reporters of the Golgi apparatus and the plasma membrane (see “Materials and methods” section). Embryos were then allowed to develop at 18 °C and imaged by confocal microscopy at different stages.

At early developmental stages (2 and 4 hours post-fertilization, hpf) the Golgi apparatus is present as separated elements. Between 6 and 8 hpf these elements begin to coalesce into larger Golgi structures, forming a single Golgi object per cell by 10 hpf (Figure 1). Single, centralized Golgi apparatuses were observed in embryonic cells at later developmental stages, up to the free-swimming larva, the pluteus (Figure S1A). To validate correct localization of our Golgi reporter EFGP_Giant-CT, we carried out co-microinjections of its mRNA and that of GalT_mCherry, encoding a commonly used fluorescently tagged Golgi targeting peptide from human galactosyl-transferase^9,10^. Indeed, the two fluorescent reporters co-localized to the same cellular structures, indicating that the C-terminal region of human Giantin (Giant-CT) correctly localizes to Golgi membranes (Figure S2). Of note, higher magnification imaging of embryos at stages following the observed clustering showed a morphology strongly reminiscent that of the Golgi ribbon observed in mammalian cells (Figure S1B).

**Figure 1.**
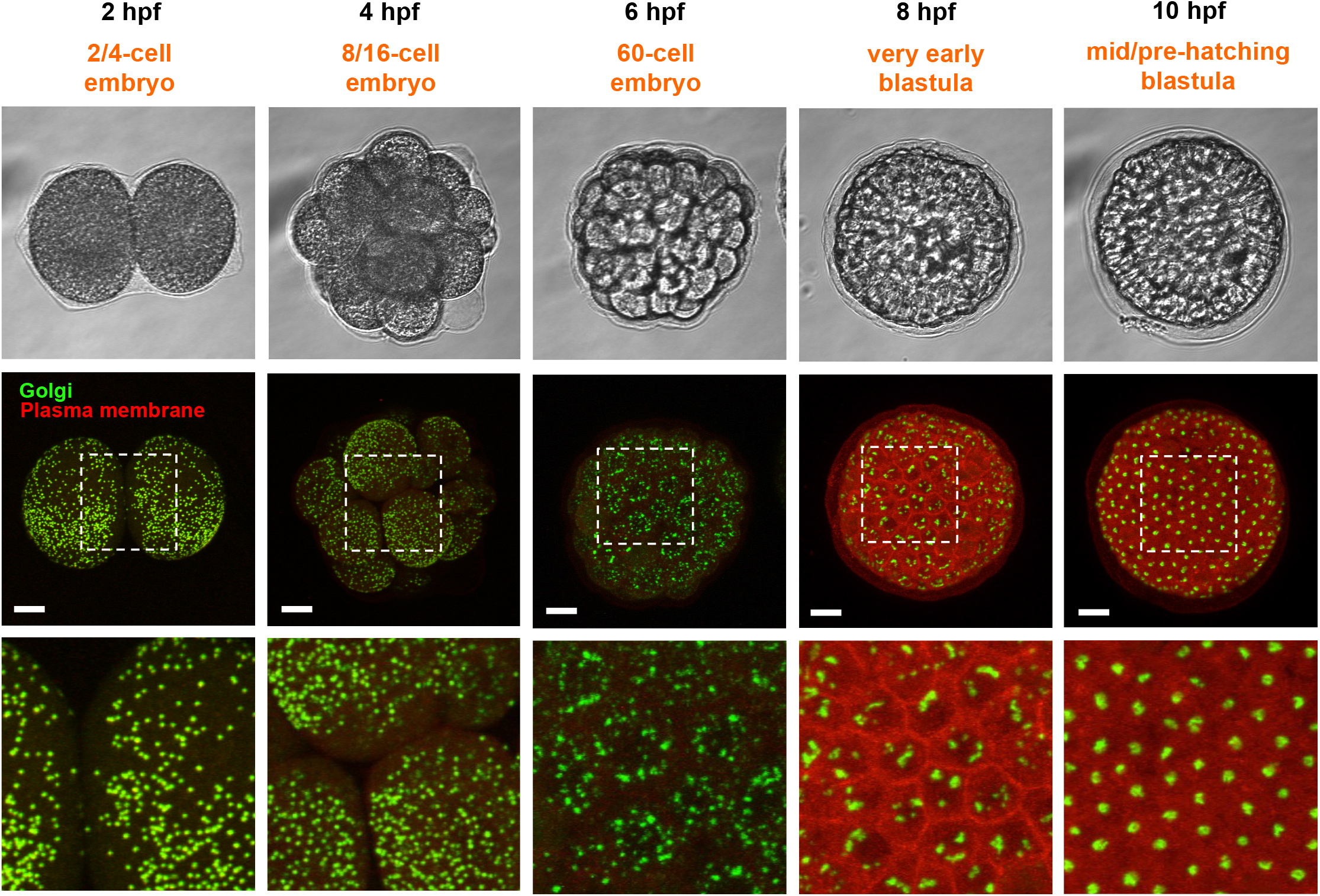
Golgi dynamics during sea urchin development. *P. lividus* zygotes were microinjected with mRNAs encoding fluorescent reporters of the Golgi apparatus (EGFP_Giant-CT) and of the plasma membrane (mCherry_CAAX) and allowed to develop at 18 °C. The indicated stages were imaged by bright field and confocal microscopy. Maximum intensity projections of image stacks are shown; bottom panels show magnifications of the middle panel insets. Scale bars: 20 µm.

Quantification of the size of Golgi elements during early development (2 to 10 hpf) confirmed clustering of Golgi elements, while the number of Golgi elements per embryo diplayed a drastic reduction at 8 hpf (Figure 2). Time-lapse confocal microscopy showed that small Golgi elements gradually clustered into larger structures, ultimately forming a single Golgi apparatus per cell at 8.30 to 9.30 hpf (Figure 3). Interestingly, Golgi elements appeared to be disassembled in cells undergoing mitosis, similar to what observed in mammalian cells^11,12^ (Figure 3).

**Figure 2.**
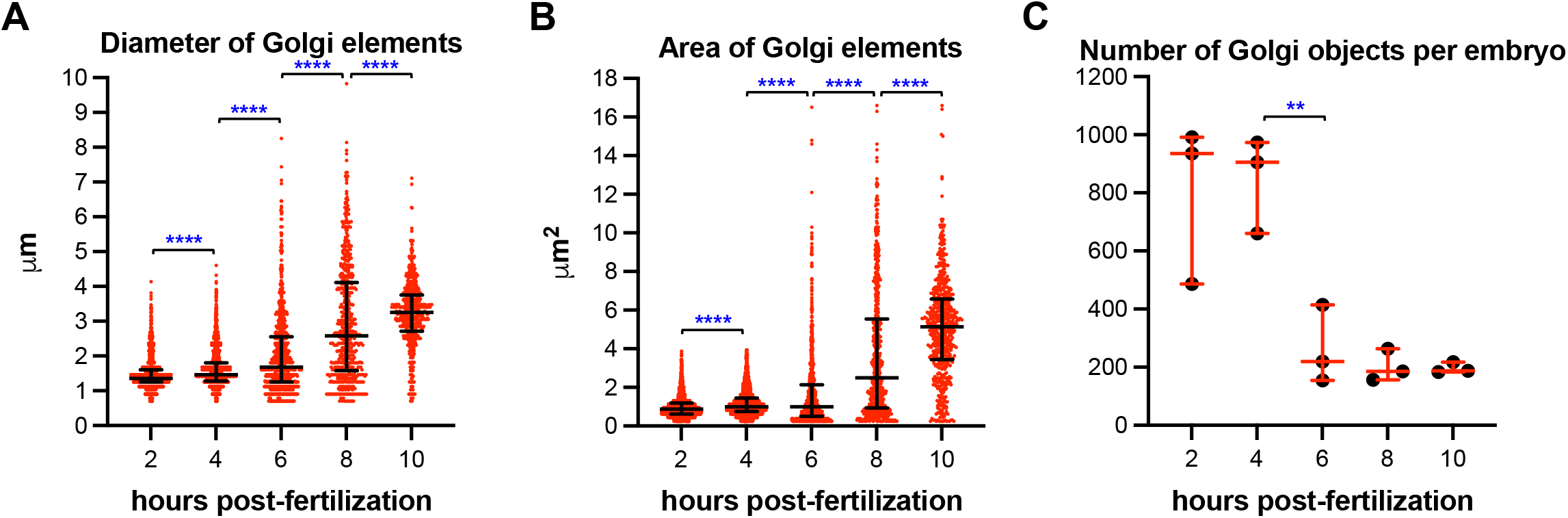
Golgi element clustering during early sea urchin development. The feret diameter (see “Materials and methods”), (A) and the area (B) of Golgi elements were quantified from three embryos per time point at 2 (N = 2274), 4 (N = 2538), 6 (N = 787), 8 (N = 604) and 10 hpf (N = 587); ****, *p* < 0.0001, Kolmogorov-Smirnov test. (C) Number of Golgi elements imaged per embryo; note that only part of the embryos could be imaged (see text) and these numbers thus underestimate the total number of Golgi objects; **, *p* < 0.01, unpaired Student’s t-test.

**Figure 3.**
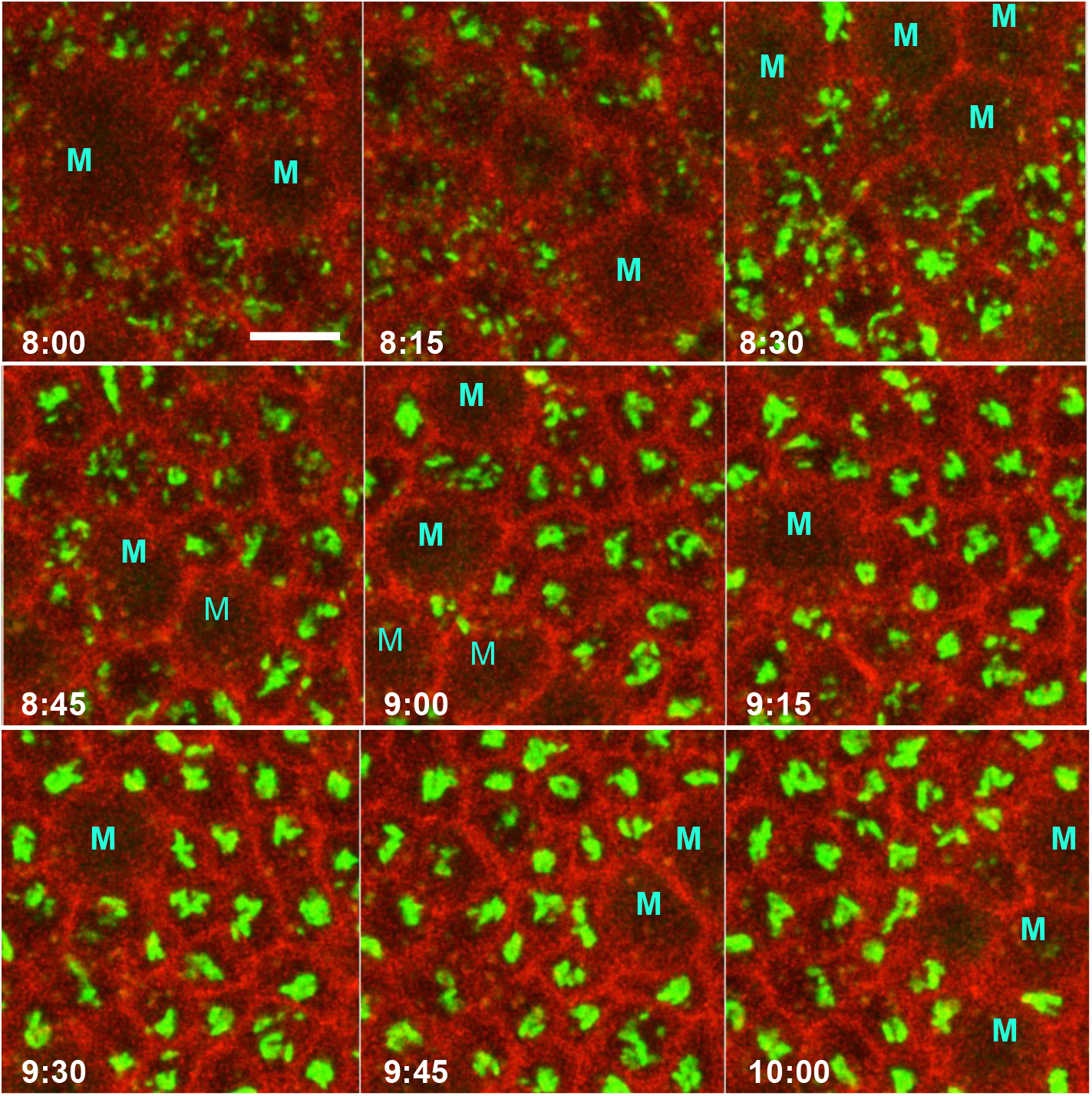
Time-lapse microscopy of Golgi clustering. mRNAs encoding reporters of the Golgi (green) and plasma membrane (red), were micro-injected in *P. lividus* zygotes, which were imaged by time-lapse microscopy at the indicated times (hpf). Note that in larger cells (labeled with M), likely undergoing mitosis, Golgi elements were barely visible, resembling the dynamics of mitotic Golgi disassembly described in mammalian cells. Scale bar: 20 µm.

While our confocal microscopy results were suggestive of ribbon formation in sea urchin embryos, we could not rule out that the observed centralized Golgi apparatuses are the result of clustering of mini-stacks but not their physical connection as in mammalian cells^13-15^. In order to ascertain that adjacent mini-stacks do connect to each other forming a true Golgi ribbon, sea urchin embryos were analyzed by electron microscopy. At 2 hpf, the Golgi was present as separate mini-stacks, whereas at 10 hpf mini-stacks were linked to each other in a typical ribbon arrangement (Figure 4). Electron microscopy thus confirms that early in development, the Golgi apparatus organization of the sea urchin switches from the typical invertebrate arrangement with separate mini-stacks to the vertebrate-like ribbon architecture.

**Figure 4.**
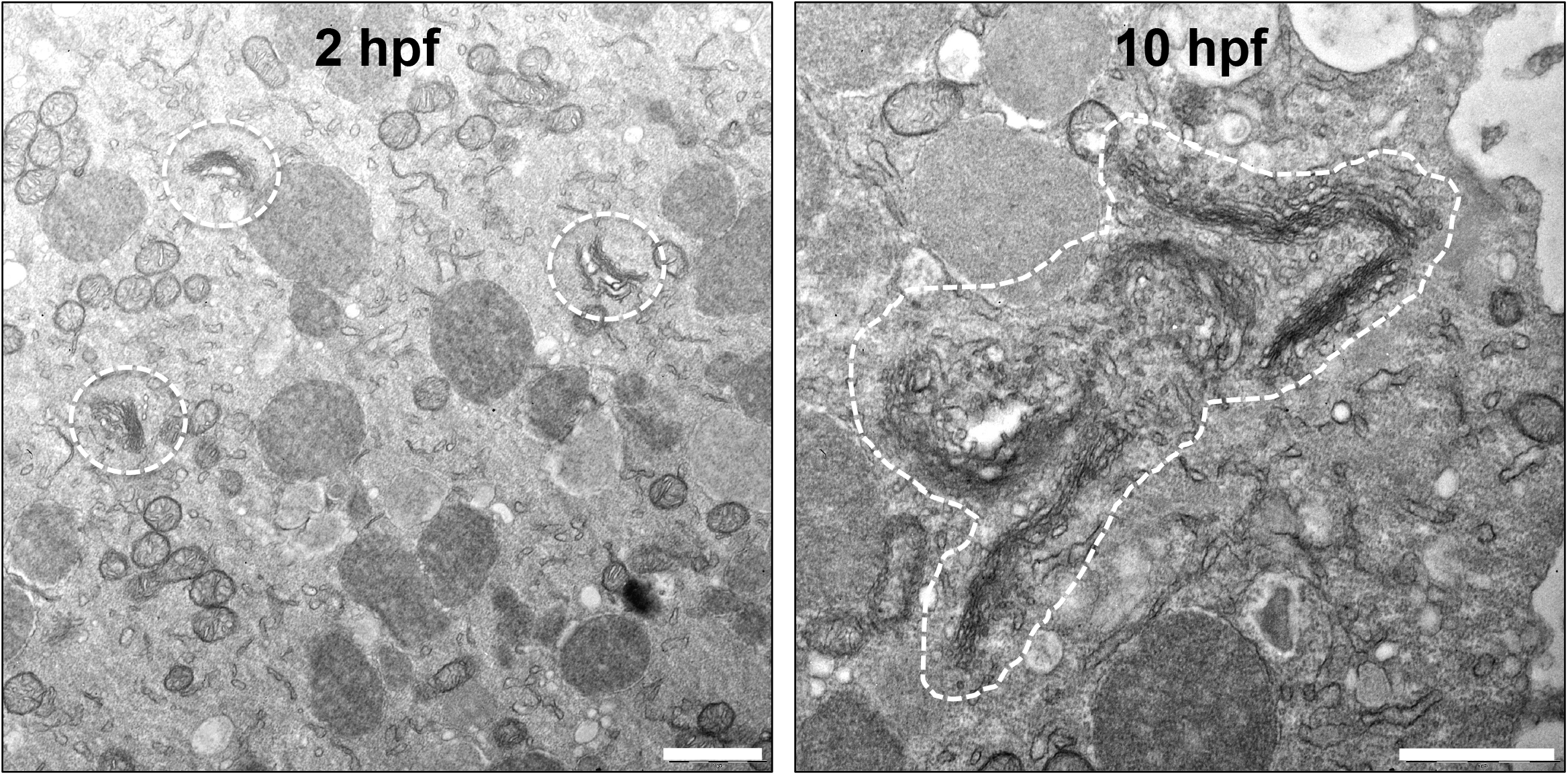
Changes in Golgi structure during *P. lividus* early development. Embryos at the indicated developmental stages were processed for electron microscopy. Golgi elements are indicated by dashed contours. Scale bars: 1 µm.

## Discussion

The Golgi apparatus plays a central role in the processing and sorting of secretory cargoes. While this biosynthetic function remains the most actively investigated^2,3,16^, recently published evidence shows that the Golgi actively participates in a number of secretion-independent cellular processes, such as stress sensing and signaling, apoptosis, autophagy and innate immunity^17-28^. The Golgi’s functional unit is the mini-stack, thus named because it is formed by a pile of flat membrane cisternae. Mini-stacks display polarization of the machinery necessary to the processing and traffic of cargo molecules from the *cis*-to the *trans*-side: the Golgi entry and exit sites, respectively. Phylogenetic analysis shows that domain functionalization within mini-stacks was already present in the last eukaryotic common ancestor^29^. Two different structural organizations of the Golgi apparatus have been described in animals. In invertebrates, the Golgi is a multi-copy organelle with separate mini-stacks dispersed throughout the cell cytoplasm; an organization seen also in plants^6,7^. In vertebrates, instead, the mini-stacks coalesce into a centralized structure, referred to as the Golgi “ribbon” after Camillo Golgi’s description, due to its appearance in optical microscopy. To date, the biological roles of the Golgi ribbon remain essentially unclear^30-32^. However, proliferating cells disassemble and reassemble the ribbon at each cell cycle, in a precisely timed and metabolically expensive process^11,33^; this level of regulation indicates that the ribbon architecture was evolutionarily selected and therefore must be functionally important. This conclusion is supported by the existence of pathologies in which ribbon breakdown (often referred to as Golgi “fragmentation”) is a hallmark^20^. Most notable among these are neurodegenerative diseases. For instance, in animal models of Amyotrophic Lateral Sclerosis (ALS), Golgi fragmentation precedes phenotypic manifestations, and in cellular models of Alzheimer’s it promotes Aβ peptide production^34-36^. Based on evidence from cultured mammalian cells, the Golgi ribbon has been proposed to mediate a variety of exocytic functions. Among these are polarized secretion, directional cell migration, Golgi enzyme homogeneity, secretion of large cargos and production of Weibel-Palade bodies, large endothelial secretory granules^37-41^. However, these hypothesized functions of the ribbon often have not been confirmed by further analyses. For instance, a study found the ribbon not necessary for the homogenous distribution of glycosylation enzymes across mini-stacks^42^. Also, repositioning of the Golgi ribbon towards the leading edge of migrating cells observed in 2D-cultures^37,38^ was neither observed in cells forced to migrate in an *in vitro* 3D-mimicking environment or *in vivo*^43^. Ribbon disassembly slows but does not block secretion of the large cargo collagen^40^. And, finally, the size of endothelial-specific secretory granules, the Weibel-Palade Bodies, is certainly controlled by ribbon integrity^41,44^, but this process is clearly a cell type-specific requirement that does not explain why most vertebrate cells make a ribbon. It is also worth considering that cells of invertebrate animals, plants and even unicellular eukaryotes have similar secretory requirements to vertebrate cells but do not make Golgi ribbons. Directional cell migration, for instance, is essential for developmental morphogenesis and wound healing of animals, including those with dispersed mini-stack Golgi architecture, such as flies and worms^45^. In conclusion, the biological activities so far proposed lack explanatory power and the functions that the Golgi ribbon mediates as a cellular structure remain an enigma.

Here, we report that, contrary to current consensus, the Golgi ribbon is not restricted to vertebrate cells. Morphological data published on sea urchins were suggestive that this might be the case^8,9^, prompting us to investigate Golgi dynamics in the Mediterranean sea urchin, *Paracentrotus lividus*. In *P. lividus* embryos the Golgi is initially present in the typical ‘invertebrate” arrangement, as dispersed mini-stacks. At subsequent stages, ribbon assembly gradually occurs and is completed by the pre-hatching blastula stage (at 10 hours hpf), persisting throughout later development, up to the free-swimming pluteus larva. Sea urchins belong to the phylum Echinodermata, early branching deuterostomes evolutionarily related to chordates and therefore vertebrates. Ribbon assembly during sea urchin development implies that: (a) this Golgi arrangement is ancient, having evolved at least before the common ancestor of deuterostomes, more than 0.6 gigayears ago; and (b) it must play some fundamental biological role(s), as it was conserved during evolution from sea urchins to humans. Interestingly, Golgi ribbon formation in early embryos is observed also in mammals. In mouse blastomeres, the Golgi is formed by separate elements, likely mini-stacks, that cluster by the blastocyst stage^46^. Formation of the Golgi ribbon in embryonic cells thus occurs at early developmental stages in both sea urchins and mammals and may indicate that this centralized Golgi organization plays a role during embryogenesis.

## Materials and Methods

### Animals

*P. lividus* adults were sourced from the Gulf of Naples and maintained at 18 °C in dedicated aquaria at the Stazione Zoologica Anton Dohrn. Gametes were obtained by vigorous shaking. For each experiment, gametes were collected from 2 to 3 males and females. Efficient fertilization was tested before proceeding to microinjections.

### Constructs

Primers were designed using the NEBuilder tool (http://nebuilder.neb.com/). PCR reactions for amplicon generation were carried out with Q5 High-Fidelity DNA Polymerase (NEB, cat. no. M0491). *pCineo_EGFP_Giant_CT*. The plasmid encodes EGFP in frame with a linker sequence (GGGSGGGS) and the 69 C-terminal amino acids of human Giantin, which target the recombinant protein to Golgi membranes^47^. The EGFP coding sequence was amplified from pEGFP-N1 vector (Clontech) with the following primers: fwd, atacgactcactataggctagcATGGTGAGCAAGGGCGAG (lower case: pCineo sequence; upper case EGFP coding sequence); rev, acctgatccaccgccCTTGTACAGCTCGTCCATGC (lower case: GGGS coding sequence; upper case: EGFP coding sequence). The sequence encoding the 69 C-terminal amino acids of human Giantin was amplified from human umbilical vein endothelial cell (HUVEC) cDNA with the following primers: fwd, *ctgtacaag*ggcggtggatcaggtggaggatctACTCCTATCATTGGCTC (italics: EGFP coding sequence; lower case: GGGSGGGS linker coding sequence; upper case: Giantin coding sequence); rev, gaggtaccacgcgtgaatTCATTACTATAGATGGCCC (lower case: pCineo sequence; upper case: Giantin coding sequence and two stop codons). *pCineo_GalT_mCherry*. A plasmid (the generous gift of Irina Kaverina, Vanderbilt School of Medicine) encoding the N-terminal 87 amino acids of galactosyl-transferase (GalT), which confer Golgi localization, in frame with mCherry^10^ was used to amplify the GalT_mCherry coding sequence using the following primers: fwd, ttaatacgactcactataggctagcATGAGGCTTCGGGAGCCG (lower case: pCineo sequence; upper case: GatT coding sequence); rev, ctctagaggtaccacgcgtgaattcTTACTTGTACAGCTCGTCCATGC (lower case: pCineo sequence; upper case: GatT coding sequence). *pCineo_mCherry_CAAX*. The sequence encoding mCherry in frame with the polybasic sequence and CAAX motif of human K-Ras (GKKKKKKSKTKCVIM) for targeting to the plasma membrane was generated by amplification of mCherry using the pmCherry-N1 (Clontech) plasmid as template and the following primers: fwd, ttaatacgactcactataggctagcATGGTGAGCAAGGGCGAG (lower case: pCineo sequence; upper case: mCherry coding sequence); rev, ctctagaggtaccacgcgtg*aattcttacataattacacactttgtctttgacttctttttcttctttttacc*CTTGT ACAGCTCGTCCATGC (lower case: pCineo sequence; italics: polybasic plus CAAX motif and stop codon coding sequence; upper case: mCherry coding sequence). Amplicons and pCineo plasmid (linearized by NheI/EcoRI digestion) were assembled using the NEBuilder HiFi DNA assembly cloning kit (New England Biolabs, cat. no. E5520), following the manufacturer instructions. Correct sequences were verified by Sanger sequencing.

### *In vitro* transcription

Plasmids were linearized by digestion with NotI, a unique restriction site in the pCineo vector located downstream of the cloned sequences. One microgram of each linearized plasmid was used as template for *in vitro* transcription, using the mMESSAGE mMACHINE T7 transcription kit (Invitrogen, cat. No. AM1344). Purified mRNAs were resuspended in DEPC-MilliQ water, their concentration measured and their quality checked by agarose gel electrophoresis. mRNAs were aliquoted and stored at – 80 °C until used.

### Microinjections

Eggs’ jelly coat was eliminated by a short wash in acidic filtered sea water (1.5 mM citric acid in 0.22 µm filtered sea water, FSW). De-jellied eggs were then immobilized on 60 mm plastic dish lids pre-treated with 1% protamine sulphate (Sigma-Aldrich, P4380) in FSW. Eggs were then washed with FSW containing sodium para-amino benzoate (Sigma-Aldrich, A6928; 0.05% in FSW) to prevent hardening of the fertilization envelope. *In vitro* transcribed mRNAs were diluted to a final concentration of 300-500 ng/µl in 120 mM KCl/DEPC-water. Four to five pl of diluted mRNAs were injected per embryo, immediately after fertilization. Embryos were allowed to develop at 18 °C.

### Confocal microscopy

At the indicated times post-fertilization, embryo development was stopped by incubation with 0.2% paraformaldehyde in FSW. This treatment kills embryos preserving EGFP and mCherry fluorescence. Since embryos were not properly fixed, imaging was carried out within 16 h of formaldehyde treatment. Embryos laid on bottom coverslip dishes containing FSW were imaged with an inverted 25x (NA 0.8) water immersion objective, using a Zeiss LSM700 system. Image stacks (z-step 1 µm) were acquired. Only one third to one half of the embryo volumes could be imaged at early stages, likely due to the opacity of yolk granules. At later stages (prism and pluteus) embryos were transparent and their whole volume could be imaged. For live imaging experiments, eggs were laid in FWS containing bottom coverslip dishes pre-treated with protamine, fertilized and then immediately microinjected with fluorescent reporter encoding mRNAs. Imaging was carried out as described above. Image stacks (z-step 1 µm) were acquired at 15 min intervals. Higher magnification imaging of embryos was carried out on EGFP_Giant-CT microinjected embryos using a 40x (NA 1.10) water immersion objective with a Leica SP8 confocal system. For presentation purposes, contrast-enhancement and Gaussian-blur filtering were carried out (ImageJ) to the images shown.

### Image analysis

Images were opened as max_int projections in ImageJ (https://imagej.nih.gov/ij/). The Golgi channel (8-bit) was selected and processed as follows.

For 2 and 4 hpf embryos: 1) background subtraction (rolling ball method, set at 50); 2) background-subtracted images were duplicated; 3) one of the images was processed to find maxima (parameter adjusted for each image to identify the majority of Golgi objects); the output is segmented particles; 4) the other image was subjected to thresholding (the value was adjusted to match Golgi object size) and then transformed into a binary image (binary>make binary); 5) the image output from step 3 was subjected to selection>create selection> copy; 6) the copied selection was pasted on binary image (step 4), then undo; 7) the pasted selection was then drawn (edit>draw) in order to separate the objects in the binary image, based on the segmentation; 8) edit>selection>select none, in order to eliminate the selection in the binary image; 9) objects in the segmented binary image were then counted and quantified (“analyze particles”; the area range was 0.25-4 µm^2^, based on tests of particle size).

For embryos from 6 hpf on: 1) open images as max_int projections of the Golgi channel (8-bit); 2) background subtraction (rolling ball method, set at 50); 3) threshold was set (best value to fit particle size) and then the image transformed from 8-bit into a binary (binary> make binary); 4) quantitation of the objects was then carried out (“analyze particles”, area range 0.25-infinite µm^2^). The “analyze particles” command generates tables with numerical values related to the objects analyzed, including their “feret diameter’, which is the longest distance between two points within an object, and their area. Graphs and statistical analyses were generated with Prism version 9 (Graphpad).

### Electron microscopy

Sea urchin embryos maintained at 18 °C were collected at the indicated developmental stages and fixed at 4° C in 2% glutaraldehyde in FSW. After 24 h samples were first rinsed in FSW (6x 10 min), then in MilliQ water (3x 10 min) and post fixed with 1% osmium tetroxide and 1.5% potassium ferrocyanide for 1h at 4° C. Samples were then rinsed five times with MilliQ, dehydrated in a graded ethanol series, further substituted by propylene oxide and embedded in Epon 812 (TAAB, TAAB Laboratories Equipment Ltd, Berkshire, UK) at room temperature for 1 d and polymerized at 60 °C for 2 d. Resin blocks were sectioned with a Ultracut UCT ultramicrotome (Leica, Vienna, Austria). Sections were placed on nickel grids and observed with a Zeiss LEO 912AB TEM (Zeiss, Oberkochen, Germany).

## Author contributions

FF designed the study. FF, GB and MIA carried out experiments. FF wrote the manuscript.

## Acknowledgements

This work was supported by Stazione Zoologica Anton Dohrn’s intramural funding to FF.

---

**Figure S1.**
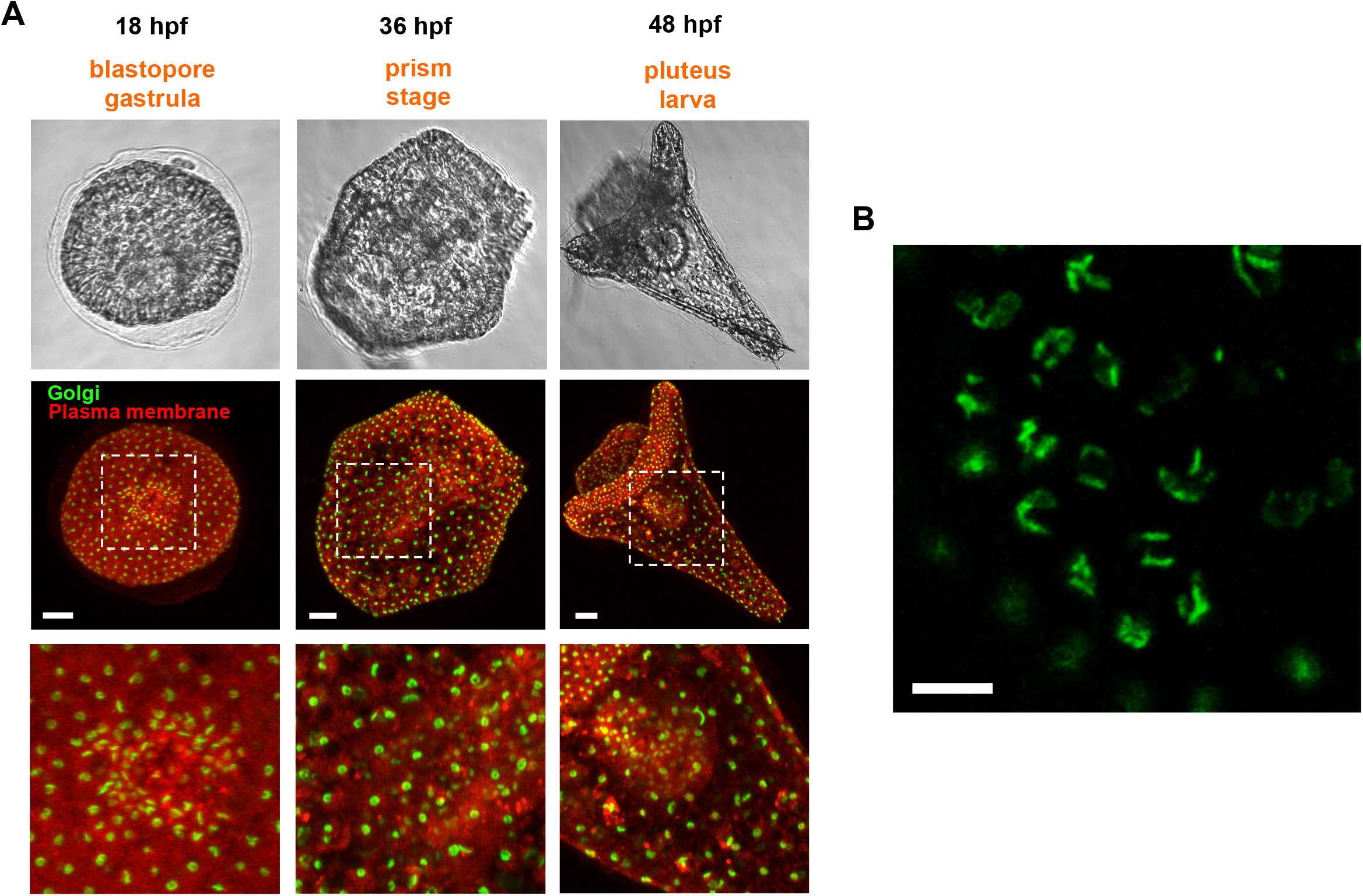
Golgi dynamics during sea urchin development. (A) *P. lividus* embryos treated as described in Figure 1 and imaged at the indicated developmental stages. Scale bars: 20 µm. (B) The Golgi apparatus in a 15 hpf *P. lividus* embryo. A single focal plane (pinhole size: 77 µm) acquired with a 40x water immersion objective is shown. Scale bar: 5 µm

**Figure S2.**
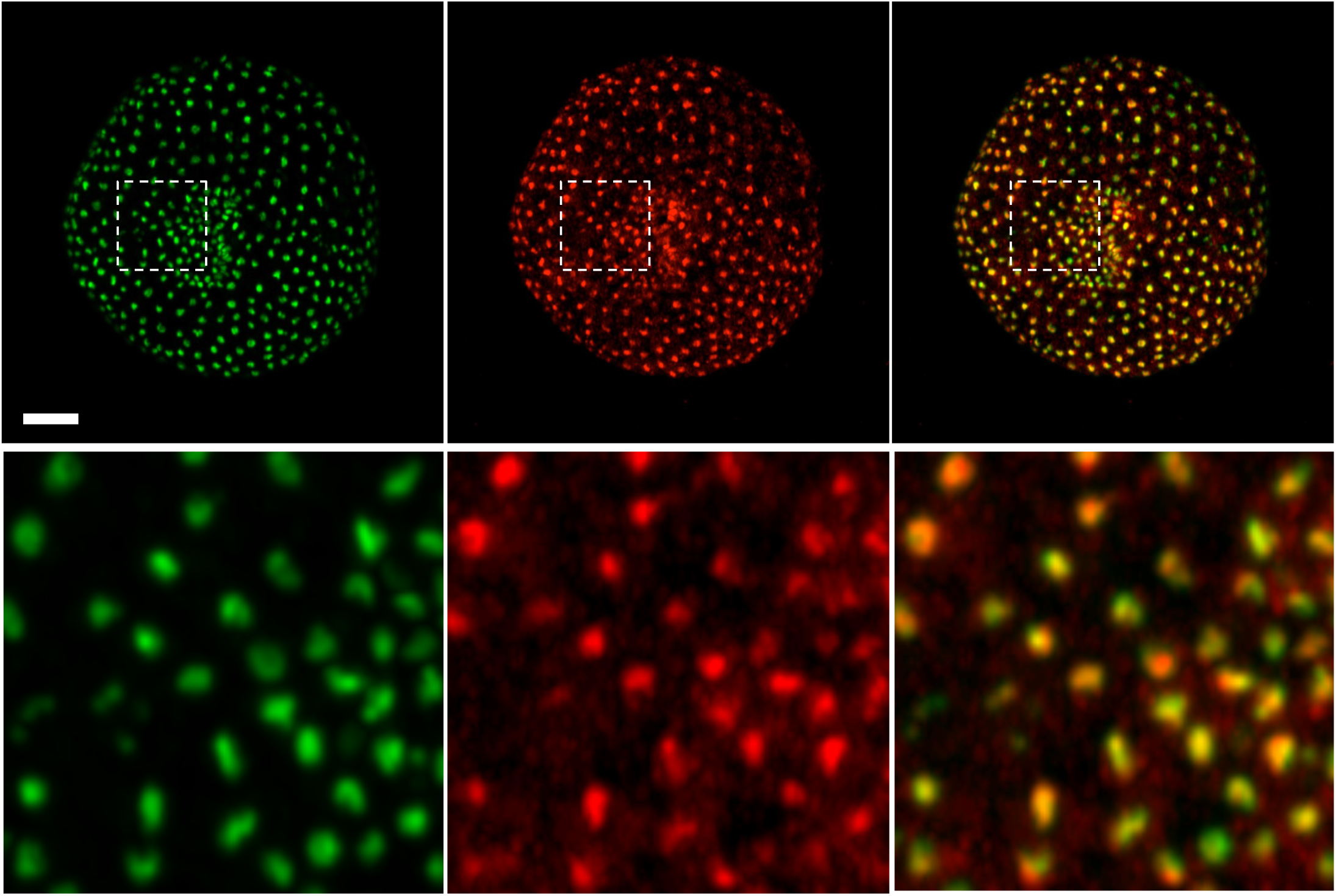
validation of the EGFP-Giant-CT reporter. EGFP_Giant-CT and GalT_mCherry encoding mRNAs were co-injected in *P-lividus* zygotes. Embryos were imaged at 21 hpf. The EGFP_Giant-CT reporter co-localizes with the established GalT reporter, indicating its correct targeting to Golgi membranes. Maximum intensity projections, acquired as described in Figure 1, are shown. Bottom panels: magnifications of the upper panel insets. Scale bar: 20 µm.

